# Phylogenomics clarifies biogeographic and evolutionary history, and conservation status of West Indian tremblers and thrashers (Aves: Mimidae)

**DOI:** 10.1101/540658

**Authors:** Jeffrey M. DaCosta, Matthew J. Miller, Jennifer L. Mortensen, J. Michael Reed, Robert L. Curry, Michael D. Sorenson

**Affiliations:** Department of Biology, Boston University, Boston, Massachusetts, USA; Biology Department, Boston College, Chestnut Hill, Massachusetts, USA; Department of Biology, Villanova University, Villanova, PA, USA; Sam Noble Oklahoma Museum of Natural History and Department of Biology, University of Oklahoma, Norman, OK, USA; Department of Biology, Tufts University, Medford, Massachusetts, USA; Department of Biological Sciences, University of Arkansas, Fayetteville, AR, USA

**Keywords:** ddRAD-seq, avian radiation, archipelago, Lesser Antilles

## Abstract

The West Indian avifauna has provided fundamental insights into island biogeography, taxon cycles, and the evolution of avian behavior. Our interpretations, however, rely on robust hypotheses of evolutionary relationships and consistent conclusions about taxonomic status in groups with many endemic island populations. Here we present a phylogenetic study of the West Indian thrashers, tremblers, and allies, an assemblage of at least 5 species found on 29 islands, which is considered the archipelago’s only avian radiation. We improve on previous phylogenetic studies of this group by using double-digest restriction site-associated DNA sequencing (ddRAD-seq) to broadly sample loci scattered across the nuclear genome. A variety of analyses, based on either nucleotide variation in 2,223 loci that were recovered in all samples or on 13,282 loci that were confidently scored as present or absent in all samples, converged on a single well-supported phylogenetic hypothesis. In contrast to previous studies, we found that the resident West Indian taxa form a monophyletic group, exclusive of the Neotropical–Nearctic migratory Gray Catbird *Dumetella carolinensis*, which breeds in North America. Earlier studies indicated that the Gray Catbird was nested within a clade of island resident species. Instead, our findings imply a single colonization of the West Indies without the need to invoke a subsequent ‘reverse colonization’ of the mainland by West Indian taxa. Furthermore, our study is the first to sample both endemic subspecies of the endangered White-breasted Thrasher *Ramphocinclus brachyurus*. We find that these subspecies have a long history of evolutionary independence with no evidence of gene flow, and are as genetically divergent from each other as other genera in the group. These findings support recognition of *R. brachyurus* (restricted to Martinique) and the Saint Lucia Thrasher *R. sanctaeluciae* as two distinct, single-island endemic species, and indicate the need to re-evaluate conservation plans for these taxa. Our results demonstrate the utility of phylogenomic datasets for generating robust systematic hypotheses.

## 1. Introduction

Island archipelagos have long been a focus for studies of biogeography and community ecology due to the potential for repeated bouts of colonization and isolation among islands, and their consequent effects on diversification rates and community assemblages (e.g., Darwin, 1859; Wallace 1869; Mayr, 1942; MacArthur and Wilson, 1967; Diamond, 1975; Grant, 1998). The spotlight of many of such studies is the West Indies, which serves as a “laboratory” for evolution and ecology due to intermediate levels of geographic isolation that allow for the accumulation of endemic lineages along with gene flow and dispersal both within the archipelago and between islands and the mainland (Ricklefs and Bermingham, 2008). The West Indies avifauna in particular has been the focus of intense study because of its species diversity, ecological diversity, and perceived capacity for dispersal. For example, the West Indies avifauna has been the subject of influential studies of colonization-extinction dynamics (Ricklefs and Birmingham, 2001), species-area relationships (Lack, 1976), and taxon cycles (Ricklefs and Cox, 1972; Ricklefs and Bermingham, 2002).

Approximately 100 bird species are endemic to the West Indies (del Hoyo et al., 2018), reflecting a complex and stochastic biogeographical history. Despite their ability to fly and presumed capacity for dispersal, the richness of the Lesser Antillean avifauna is less than predicted by equilibrium models given the number of potential colonists from North and South America (Ricklefs and Birmingham, 2001), suggesting that expanses of open water between the mainland and the islands represent a significant barrier for many birds. Factors such as heterogeneous patterns of colonization and extinction (Ricklefs and Birmingham, 2001), the geography of colonization sources (Cox and Ricklefs, 1977), ecological constraints (Faaborg, 1985), and host-pathogen dynamics (Fallon et al., 2003) are also thought to influence the richness of different clades. Recent human activities have also played a role in shaping the West Indian avifauna through introductions (e.g., Ricklefs and Birmingham, 2008), changes in land use (Acevedo and Restrepo, 2008; Young et al., 2010), and their effects on extinction risk (Williams and Steadman, 2001).

The West Indies avifauna lacks the remarkable adaptive radiations seen in the Galapagos or Hawaiian Islands, presumably due to a relative dearth of available niche space resulting from higher rates of colonization (Ricklefs and Birmingham, 2008). One exception, however, is a group of five species of thrashers and tremblers (family Mimidae) that are year-round residents of the West Indies. Two of these species (White-breasted Thrasher *Ramphocinclus brachyurus* and Gray Trembler *Cinclocerthia gutturalis*) are both endemic to the Lesser Antillean islands of Saint Lucia and Martinique, whereas the remaining species are widespread across the Lesser Antilles (Scaly-breasted Thrasher *Allenia fusca* and Brown Trembler *Cinclocerthia ruficauda*) or the Lesser Antilles, Puerto Rico, Bahamas, and Bonaire (Pearly-eyed Thrasher *Margarops fuscatus*) (Cody, 2005). Previous phylogenetic analyses of these species have been based on data from the mitochondrial genome and 1–4 nuclear loci (Hunt et al., 2001; Lovette and Rubenstein, 2007; Lovette et al., 2012; Figure 1). These studies revealed considerable mitochondrial genetic divergence among genera and within *Cinclocerthia* consistent with long histories of isolation among these taxa. These West Indian species are members of a clade that also includes the ‘blue’ mockingbirds of Mexico and Central America (Blue Mockingbird *Melanotis caerulescens* and Blue-and-white Mockingbird *M*. *hypoleucus*) and the catbirds of southeastern Mexico and Belize (Black Catbird *Melanoptila glabrirostris*) and North America (Gray Catbird *Dumetella carolinensis*) (Lovette et al., 2012). *Dumetella carolinensis* is the only long-distance migratory species in the clade. It breeds across much of the United States and southern Canada, but moves during the non-breeding season to continental regions adjacent to the Atlantic, including the Gulf Coast, Mexico, and Central America, as well as northwestern Colombia and the West Indies (Lucayan Archipelago and Greater Antilles) (Cody, 2005). Interestingly, mitochondrial gene trees strongly support a sister relationship between *Dumetella carolinensis* and *Ramphocinclus brachyurus*, rendering the group of resident West Indies species paraphyletic (Hunt et al., 2001; Lovette and Rubenstein, 2007; Lovette et al., 2012). This finding was hypothesized to be the result of a single colonization of the West Indies followed by ‘reverse colonization’ of an ancestor to the mainland *Dumetella carolinensis* (Lovette et al., 2012). However, gene trees based on the small number of nuclear loci that have been analyzed lack the requisite resolution to confidently place *Dumetella carolinensis* (Hunt et al., 2001; Lovette and Rubenstein, 2007; Lovette et al., 2012).

**Fig. 1.**
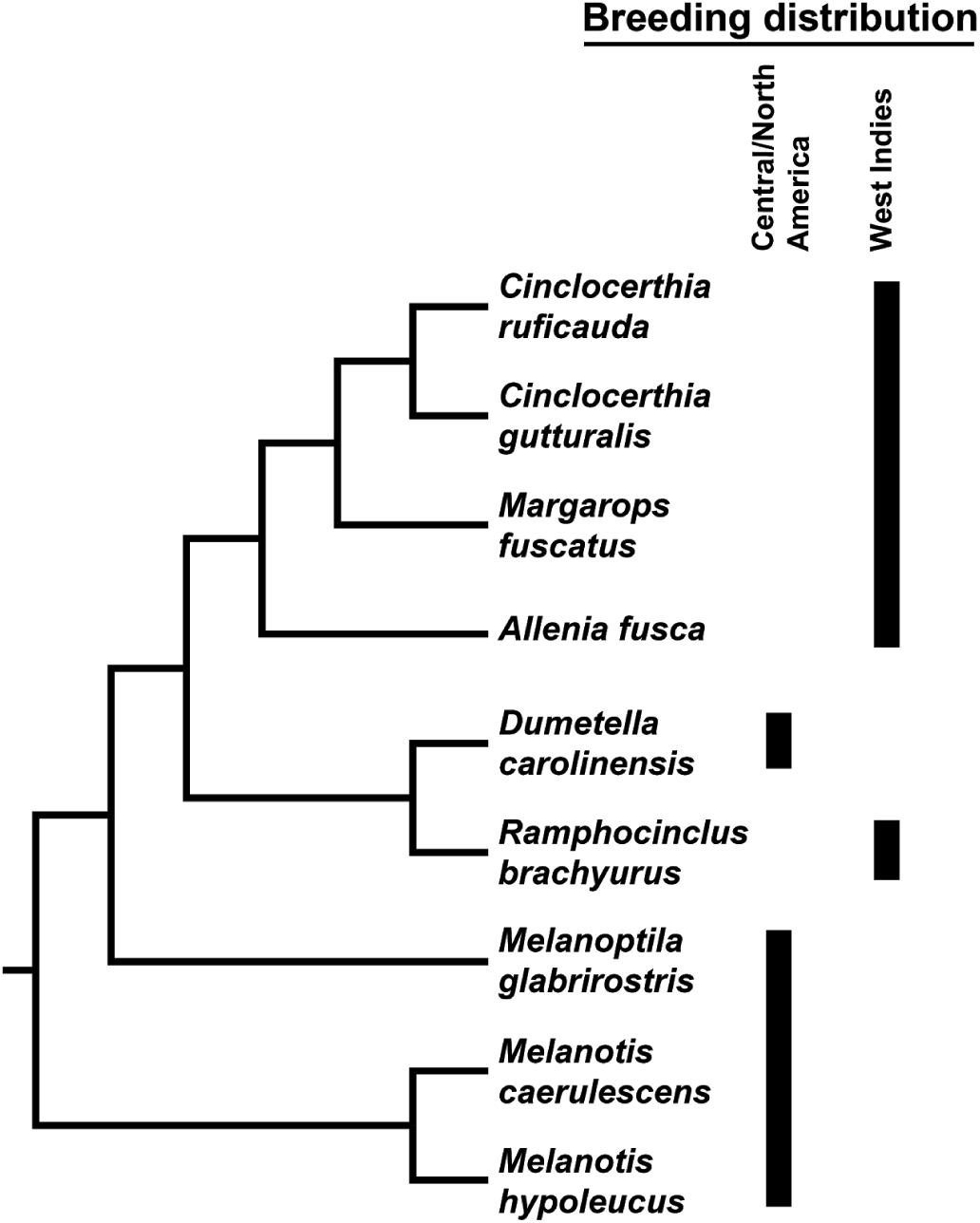
Current phylogenetic hypothesis for West Indian thrashers, tremblers, and allies. The figure is based on the analysis of Lovette et al. (2012), which used sequence data from the mitochondrial genome and three nuclear loci. All nodes had posterior probabilities > 0.95 in a partitioned Bayesian analysis using all loci.

The West Indies avifauna currently faces increased extinction pressures due to human activities such as habitat degradation through land use changes and introductions of diseases and predators (Ricklefs and Birmingham, 2008). Among the West Indian thrashers and tremblers, *Ramphocinclus brachyurus* is the only species with a sensitive conservation status (Mortensen et al., 2017); it is currently classified as endangered with a declining population trend (IUCN, 2018). This species has a narrow geographic distribution, with distinct subspecies occupying Martinique (*brachyurus*) and Saint Lucia (*sanctaeluciae*). The subspecies differ in plumage, ecology, and potentially calls (John 1995; Cody, 2005). The subspecies *sanctaeluciae*, which has been considered a separate species (Cory, 1887; Ridgway, 1907), has a larger population but has saturated available suitable habitat on the island (Sass et al., 2017). Elevating *sanctaeluciae* to species status would likely result in IUCN Red-list categories of endangered for *sanctaeluciae* and critically endangered for *brachyurus* in Martinique. No previous phylogenetic study has sampled both of these subspecies.

Here we test the current phylogenetic hypothesis of the West Indian thrashers, tremblers, and allies (Fig. 1) using double-digest restriction-site associated DNA sequencing (ddRAD-seq) to generate thousands of nuclear markers scattered across the genome. We use several data matrices and analyses to investigate: 1) the relationships among genera, particularly the phylogenetic placement of *Dumetella carolinensis*; 2) the level of genetic divergence between the endangered subspecies *R. b. brachyurus* and *R. b. sanctaeluciae*; and 3) population structure within the widespread species *Cinclocerthia ruficauda*.

## 2. Materials and Methods

### 2.1. Taxon sampling and DNA extraction

We collected tissue or blood samples for 26 mimid samples from 11 species (Table 1). Each species was represented by 1-6 samples, with an emphasis on greater sampling of species with described subspecies. Total genomic DNA was extracted from each sample using the Qiagen DNA Blood and Tissue Kit following the manufacturer’s protocol.

**Table 1.**
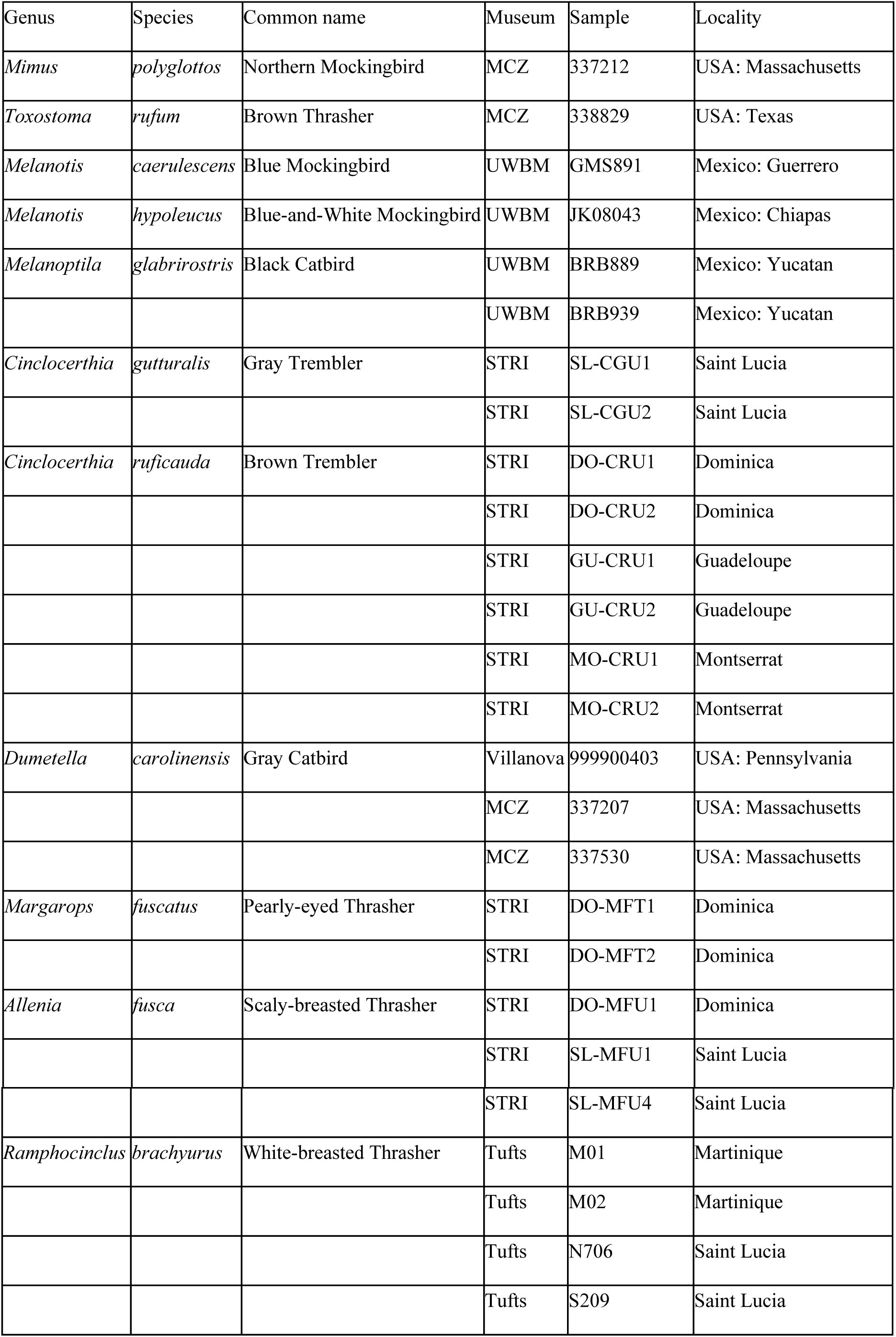
Museum and locality information for samples used in this study. MCZ: Museum of Comparative Zoology; STRI: Smithsonian Tropical Research Institute; UWBM: University of Washington Burke Museum.

### 2.2. ddRAD-seq laboratory protocols

We generated ddRAD-seq libraries for each sample following the protocol described in DaCosta and Sorenson (2014), which we briefly outline below. We double-digested total genomic DNA with the restriction enzymes SbfI and EcoRI. We then ligated barcoded adapters to the sticky ends, such that each sample received a unique adapter/barcode combination. The barcode was located inline on the SbfI-end adapter such that reads started with a 6 base pair (bp) barcode followed by a partial SbfI site; a second barcode on the EcoRI-end was located within the adapter sequence and was collected with an index read. Gel electrophoresis was used to select fragments+adapters in the 300-450 bp size range (corresponding to 178-328 bp of sample DNA), although smaller fragments were also carried over during size selection (see DaCosta and Sorenson, 2014). To reduce amplification bias toward smaller fragments, we cut products from the gel using a tapered “wedge” cut such that the full width of the gel at 450 bp and half of the gel’s width at 300 bp was retained. Fragments were then PCR amplified. The adapter associated with EcoRI (i.e., the more common cutter) included a “divergent-Y” so that fragments with EcoRI sticky ends on both sides were not amplified. We amplified products using SPRI beads, we estimated the concentration of each library via qPCR using a KAPA Biosystems kit. We pooled libraries in equimolar concentrations and 151 bp single end reads were collected from the multiplexed library using Illumina TruSeq chemistry on an Illumina HiSeq 2500 machine at the Tufts University Core Facility. De-multiplexed fastq files for all samples are available in the National Center for Biotechnology Information Short Read Archive (BioProject: PRJNA514402).

### 2.3. Bioinformatics

Sequence reads were first parsed into batches with distinct EcoRI-end adapters based on index sequences at the core facility using CASAVA v1.8 (Illumina, Inc.). We processed reads using the pipeline described in DaCosta and Sorenson (2014; see also https://github.com/BU-RAD-seq), which is briefly described below. Reads within batches were parsed to individual samples using the inline barcode sequences, which were subsequently trimmed using a custom script included in the pipeline. A “CC” was then added to the beginning of each read to reconstruct the SbfI restriction site. For short loci (i.e., those for which sequence reads reached the distal adapter), the adapter sequence was trimmed and a “C” was added to the end of the sequence to reconstruct the EcoRI restriction site. For each sample, identical reads were collapsed (with counts maintained) using the UCLUST tool in USEARCH v5 (Edgar, 2010). We removed reads with an average quality < 20 that did not cluster with any other reads. Condensed and filtered reads for all samples were concatenated into a single file, and UCLUST was used to cluster putative loci with an identity threshold of 85%. We mapped the highest quality read from each cluster to the zebra finch (*Taeniopygia guttata*) reference genome using BLAST v2.2.25 (Altschul et al., 1990). Clusters with BLAST hits to identical or nearly identical genomic locations were merged, and each cluster was then aligned using MUSCLE v3.8.31 (Edgar, 2004). We used the script RADGenotypes.py to discover and genotype polymorphisms within each locus. This script codes individual genotypes for each locus into four main categories: “missing” (no reads recovered), “good” (unambiguously genotyped), “low depth” (< 5 reads), or “flagged” (ratio of alleles divergent from 50:50 expectation or > 2 alleles). Putative loci for which multiple individual genotypes are flagged most often represent two or more paralogous loci with sufficient sequence similarity to be clustered together.

For phylogenetic analyses based on sequence data, we took a conservative approach and analyzed only loci that were variable and represented by 26 “good” genotypes (i.e., no samples with missing, low depth, or flagged genotypes) to produce a dataset of high quality, single-copy loci with no missing data. This resulted in a dataset with 2,223 loci. Because strong filters against missing data in ddRAD datasets could potentially bias phylogenetic analyses (Huang and Knowles, 2016), we also generated a data matrix of binary characters based on the presence-absence of ddRAD loci among samples/taxa. Variable recovery of loci among taxa is most often due to polymorphisms within the restriction sites, and the resulting presence-absence data has been shown to have utility in phylogenetic analyses (DaCosta and Sorenson, 2016). Scoring presence or absence of a locus can be complicated by low sequencing depth, variance among samples in size selection, and/or star activity (i.e., non-specific cutting by the restriction enzymes; DaCosta and Sorenson, 2014). Therefore, we only scored loci for which all samples had a sequencing depth of either zero or ≥ 10, which resulted in a matrix with 13,282 binary characters (loci).

### 2.4. Phylogenetic analyses

In one set of analyses, we concatenated sequences from the set of 2,223 loci (see above) into a single data matrix, with the two alleles for each sample collapsed into a single consensus sequence using IUPAC codes and an N for positions with a gap in one or both alleles. From this matrix we estimated a phylogenetic network, a maximum likelihood tree, and a species tree based on quartet inference. We generated the phylogenetic network using SPLITSTREE v4.14.6 (Huson and Bryant, 2006), employing the neighbor-net algorithm. For the maximum likelihood phylogeny, we used RAxML v8 (Stamatakis 2014) with the GTRGAMMA model of molecular evolution, 20 searches for the best topology, and 1000 standard bootstrap replicates. Based on the placement of *Dumetella* samples in the maximum likelihood tree (see Results), we also used RAxML to conduct a Shimodaira-Hasegawa (SH) test (Shimodaira and Hasegawa, 1999), comparing our best tree to a topology in which *Dumetella* and *Ramphocinclus* samples form a clade as previously hypothesized (Lovette and Rubenstein, 2007; Lovette *et al.*, 2012). The species tree was estimated using the multispecies coalescent with the SVDQuartets method (Kubatko and Chifman, 2014) in PAUP* v4.0a (Swofford, 2002) with exhaustive quartet searches and 1000 bootstrap replicates.

We also estimated species trees using biallelic polymorphisms (either SNPs or indels) as input data for the SNAPP module (Bryant et al., 2012) built into BEAST2 v2.1.2 (Bouckaert et al., 2013). One biallelic polymorphism was randomly selected from each ddRAD locus to assure independence among characters, and the analysis was repeated with a second random draw of polymorphisms. These data matrices contained 2,222 characters because for one locus the only polymorphism was a single tri-allelic SNP. For each data matrix we completed two runs with different random seed numbers, default model parameters, the two alleles from each sample assigned to a single “taxonset,” two million generations, and sampling every 5000 generations (i.e., 400 samples). Convergence within and between runs was assessed in TRACER v1.5. We trimmed the first 40 samples (10%) as burnin, and joined runs to form a posterior distribution of 720 samples.

For the presence-absence of ddRAD loci matrix (see above), we searched for the maximum parsimony phylogeny in PAUP* with all characters equally weighted, starting trees obtained through random stepwise addition with 10 replicates, the tree-bisection-reconnection swapping algorithm, and 1000 bootstrap replicates.

### 2.5 Intraspecific population structure

We found strong evidence of island monophyly for all species found on multiple West Indian islands except for *Cinclocerthia ruficauda*. Therefore, for this species, we further investigated population differentiation using STRUCTURE v2.3.4 (Pritchard et al., 2000). For input data, we subsampled the 2,223 locus dataset for loci that were variable within this species and then randomly drew one biallelic polymorphism (SNP or indel) per locus. Because the program can fail to detect population structure with ddRAD data when uninformative variants with a minor allele count of one are included (Linck & Battey, 2017), we restricted the analysis to polymorphisms with a minor allele count of two or more. This resulted in a data matrix of 643 characters. The program was run (0.5M burnin and 3M reps of MCMC) with *K*, the number of populations, set from 1 to 4. At each level of *K*, 10 replicate runs were completed and the five replicates with the highest likelihood were combined using CLUMPP v1.1.2 (Jakobsson and Rosenberg, 2007). The robustness of the results was assessed with a second set of analyses based on a different random draw of one biallelic polymorphism per locus.

## 3. Results

### 3.1 ddRAD data characteristics

The mean number of filtered reads was 995,282 per sample (stdev: 163,544), while the mean number of loci with 10 or more reads was 9,105 per sample (stdev: 186). The low variance in these measures indicates high quality data across all samples. Limiting the dataset to loci with unambiguous genotypes for all samples yielded 2,223 variable loci comprising a total of 274,456 alignment positions. These loci had an average of 12 polymorphisms (stdev: 6.6), and all but one of these loci included at least one biallelic polymorphism. Our presence-absence dataset comprised 13,282 binary characters (i.e., loci), of which 2,617 loci were present in all samples and 5,028 were parsimony informative.

### 3.2 Phylogenetic analyses

Analyses based on the concatenated 2,223 locus dataset converged on a consistent and well-supported hypothesis for the species- and genus-level relationships among these thrashers and tremblers (Fig. 2). Monophyly of each genus was supported by high bootstrap values in the maximum likelihood and SVDquartets trees and by long edges and short distances between parallel edges among genera in the phylogenetic network. Collectively, these findings provide strong evidence that each currently-recognized genus is monophyletic, including all genera breeding in the West Indies (*Cinclocerthia, Maragarops, Allenia*, and *Ramphocinclus*). Finally, rather than being embedded within a West Indian clade as found in previous studies, *Dumetella carolinensis* is unambiguously supported as the sister taxon of *Melanoptila glabrirostris*. In turn, this clade is sister to a clade comprising all the resident West Indian species. An alternative topology including a *Dumetella*–*Ramphocinclus* clade was rejected based on the SH test (p < 0.01).

**Fig. 2.**
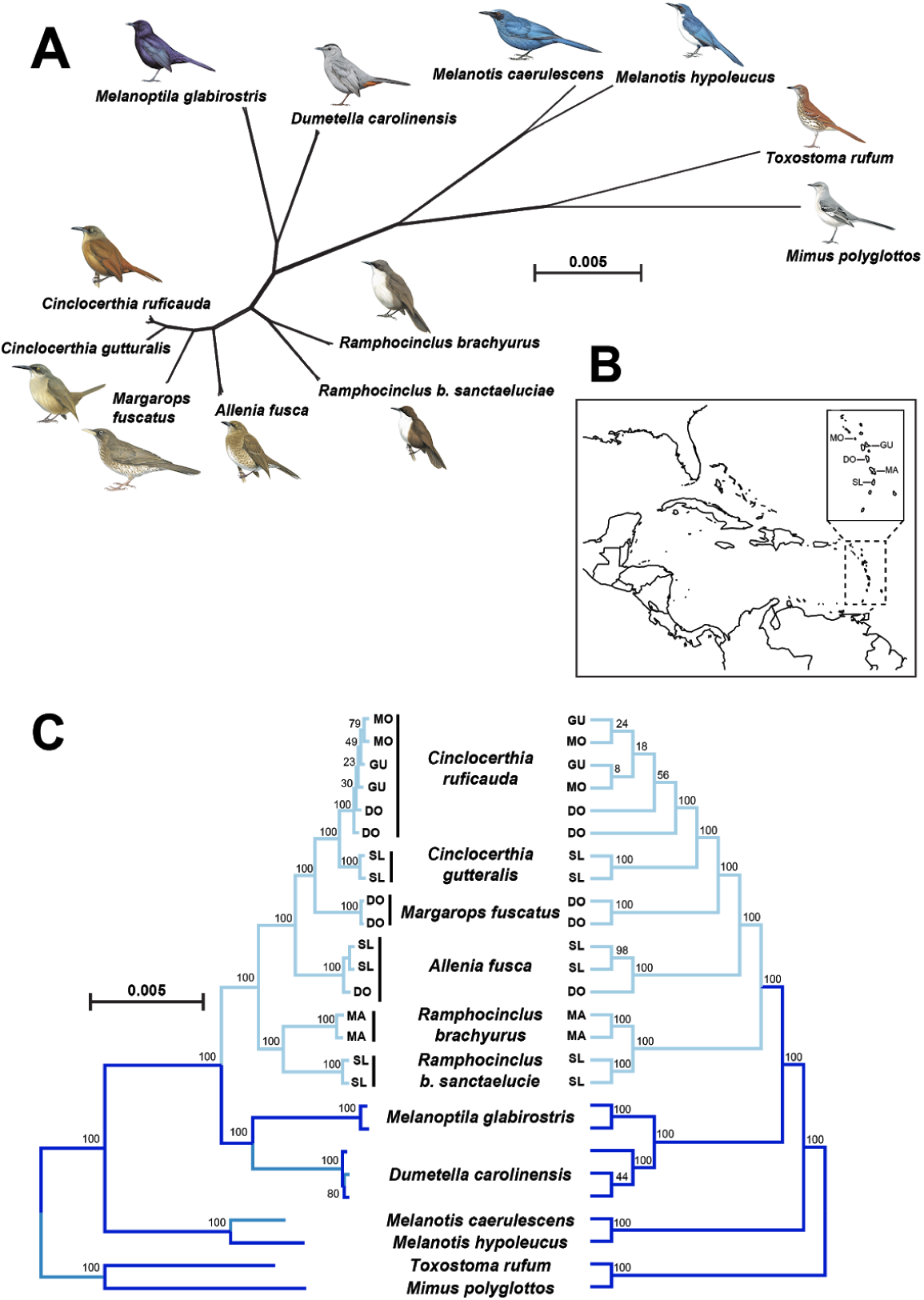
Phylogenetic hypotheses for thrashers and tremblers based on a concatenated matrix of 2,223 ddRAD loci. (A) Results for the neighbor-net phylogenetic network analysis. with bird plates from del Hoyo et al. (2018). (B) Map of the West Indies with an insert showing the islands where individuals were sampled in this study. (C) Results for concatenated maximum likelihood (left) and quartet-based species tree (right) analyses. Numbers at nodes show bootstrap percentages. From north to south, MO: Montserrat; GU: Guadeloupe; DO: Dominica; MA: Martinique; SL: Saint Lucia. Dark blue (dark gray in grayscale) branches represent taxa breeding in continental Central and North America Light blue (light gray in grayscale) branches represent all West Indian resident taxa, indicating that this radiation likely arose from a single colonization event.

The SNAPP species tree analysis based on one randomly drawn biallelic polymorphism from 2,222 loci produced consensus trees consistent with the above results, but with somewhat more uncertainty for a few interspecific relationships (Fig. 3). Separate analyses based on different random draws also produced congruent results. These analyses recovered a sister relationship between *Dumetella carolinensis* and *Melanoptila glabrirostris* as well as monophyly of the resident West Indian genera (1.0 posterior probabilities). However, relationships among *Cinclocerthia, Maragarops*, and *Allenia* were non-significant in these analyses. Alternative topologies are visualized in the cloudogram of posterior trees (Fig. 3).

**Fig. 3.**
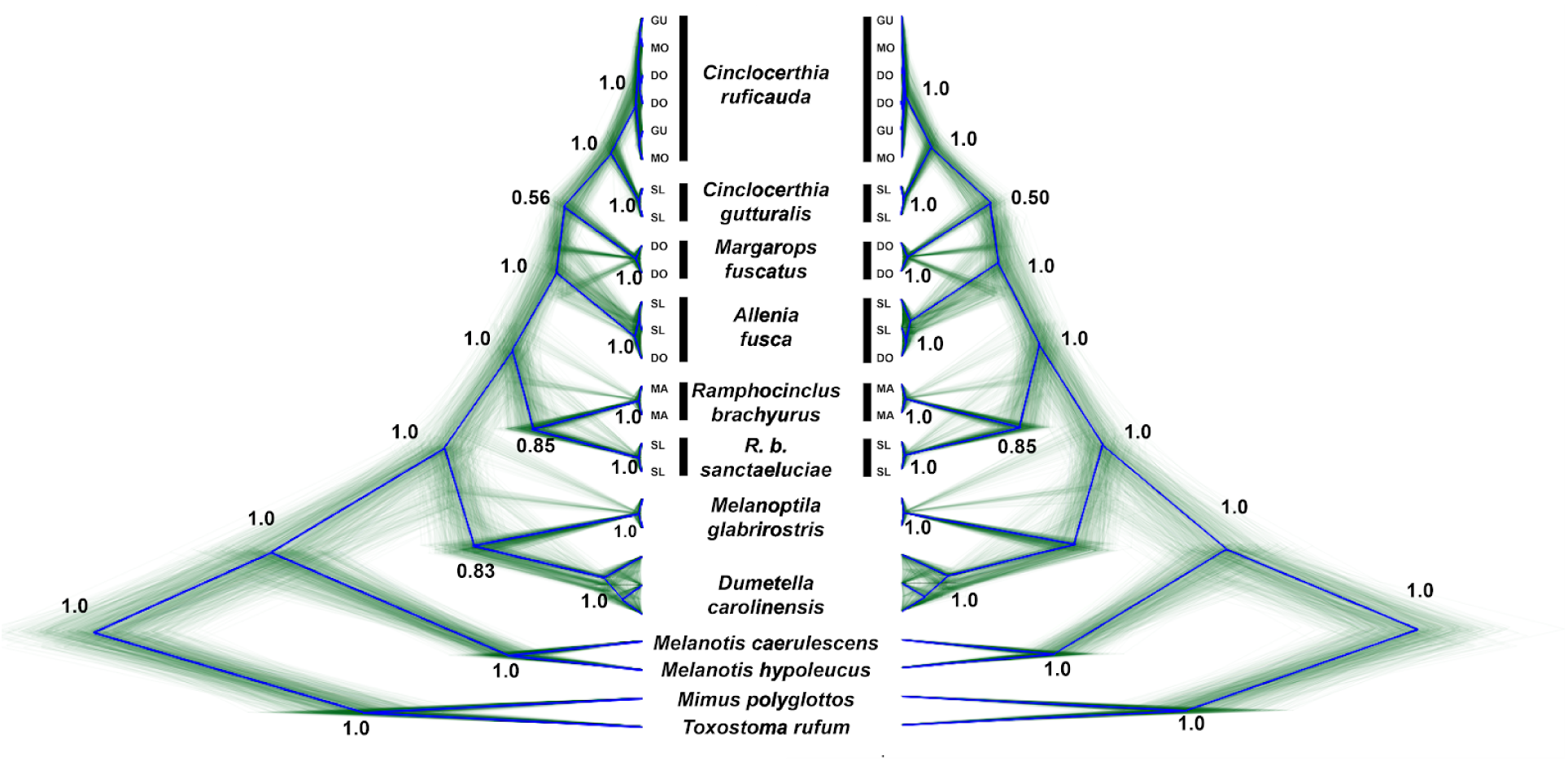
Cloudograms illustrating the posterior distributions from SNAPP species tree analyses based on one randomly drawn bi-allelic polymorphism from each of 2,222 ddRAD loci. Mirrored cloudograms show results from analyses based on two different random draws of SNPs. Green and blue lines display individual samples from the posterior distribution and the posterior consensus, respectively. Numbers at selected nodes show the posterior probability. DO: Dominica; GU: Guadeloupe; MA: Martinique; MO: Montserrat; SL: Saint Lucia.

The parsimony analysis based on presence-absence of ddRAD loci produced a phylogeny that was similar to other datasets and methods (Fig. 4), except that *Cinclocerthia ruficauda* was paraphyletic with respect to *Cinclocerthia gutturalis*, albeit with a low bootstrap value (86%) given the size of the data matrix. As in the above analyses, all taxa resident in the West Indies form a monophyletic group that is sister to the *Melanoptila–Dumetella* clade. Likewise, this analysis supports monophyly of each genus with 100 percent bootstrap values.

**Fig. 4.**
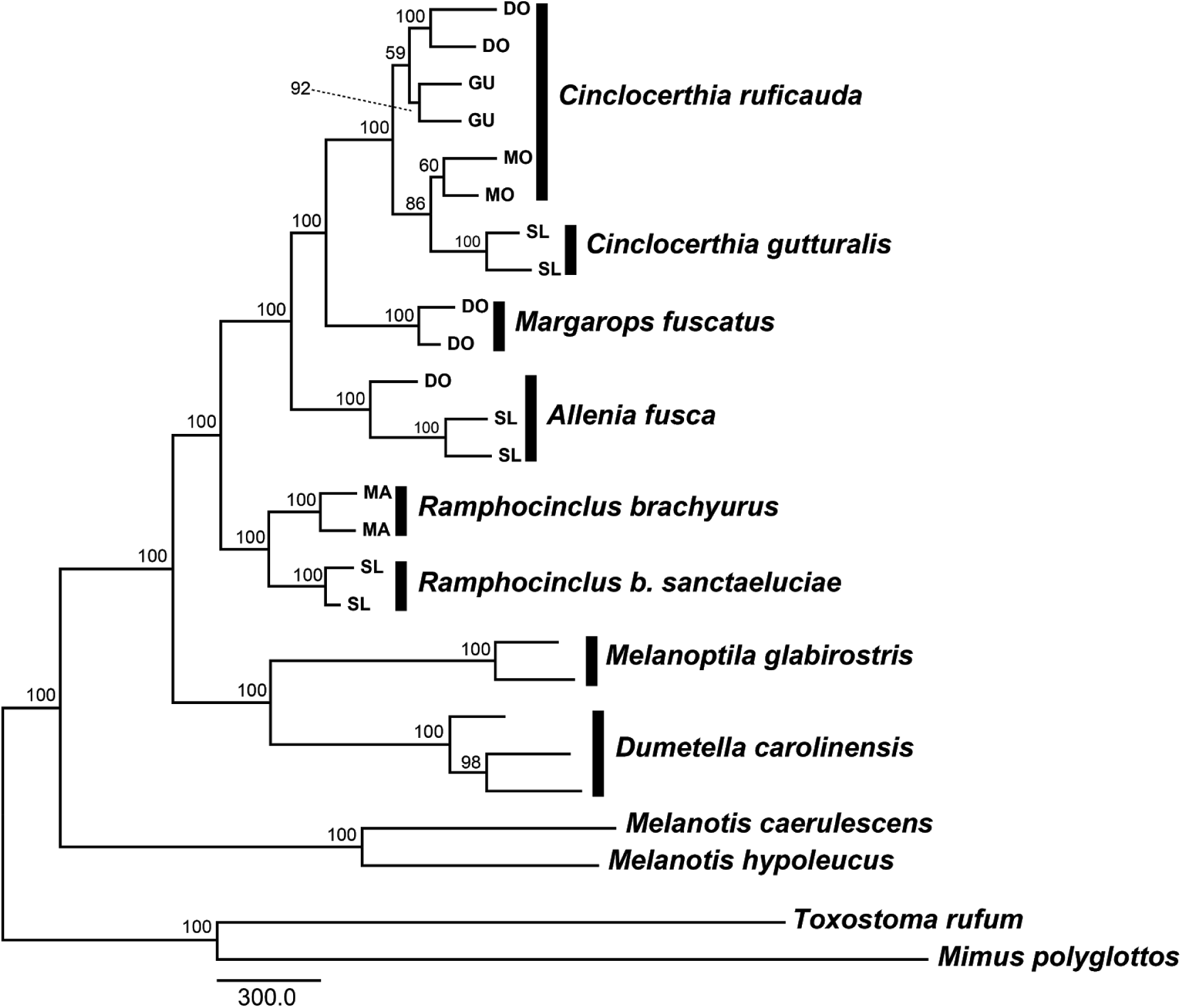
A maximum parsimony phylogeny based on the presence-absence of 13,282 ddRAD loci. Numbers at nodes show bootstrap percentages. DO: Dominica; GU: Guadeloupe; MA: Martinique; MO: Montserrat; SL: Saint Lucia.

### 3.3 Intraspecific phylogenetic variation

Our phylogenetic analysis revealed considerable intraspecific divergence for multiple species (Figs. 1-3). Most notably, samples of *Ramphocinclus brachyurus* from Martinique (subspecies *brachyurus*) and Saint Lucia (subspecies *sanctaeluciae*) are highly divergent, with branches between samples from these islands similar in length to those separating the various West Indian genera from each other. The other West Indian species sampled on multiple islands are *Allenia fusca* and *Cinclocerthia ruficauda*. For both of these species, genetic divergence among islands was more shallow than among *R. brachyurus* populations, and for *Cinclocerthia ruficauda* there was no evidence that island populations comprise monophyletic groups in our phylogenetic results (Fig. 1). However, STRUCTURE analyses of *Cinclocerthia ruficauda* support population differentiation between Montserrat and the other sampled islands (Guadeloupe and Dominica), with no further population structure evident (Fig. 5). We also recovered substantial genetic diversity among three samples of *Dumetella carolinensis*.

**Fig. 5.**
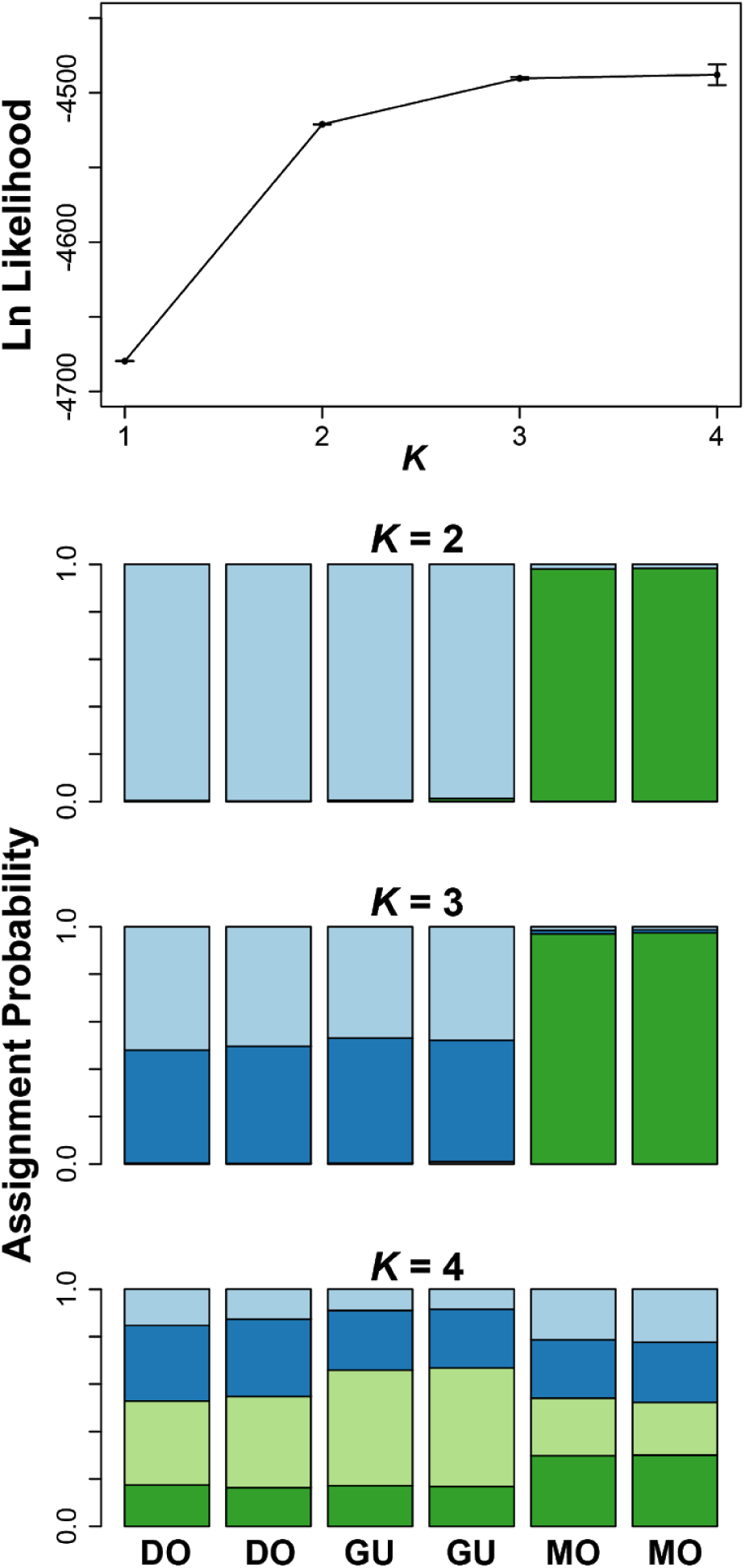
STRUCTURE results for six *Cinclocerthia ruficauda* samples. Analyses were run with one randomly drawn bi-allelic polymorphism from each of 643 ddRAD loci. DO: Dominica; GU: Guadeloupe, MO: Montserrat.

## 4. Discussion

### 4.1 Evolutionary relationships among genera and biogeographic implications

We used thousands of ddRAD loci to conduct phylogenetic analyses of the West Indian thrashers and tremblers and closely related mockingbirds and catbirds. Our study represents far wider sampling of the nuclear genome than previous molecular phylogenetic studies of these species (Hunt et al., 2001; Lovette and Rubenstein, 2007; Lovette et al., 2012). Our analyses using different methods and/or data matrices were largely consistent, and strongly supported the hypothesis that each genus is monophyletic and evolutionarily divergent from each of the other genera (Figs. 2–4). Inferred relationships are similar to those reported in previous studies, with the ‘blue’ mockingbirds (*Melanotis*) sister to all other genera, and a clade comprising *Allenia, Margarops*, and *Cinclocerthia*. Within this clade, all but the SNP-based species tree analysis (SNAPP) converged on a *Margarops–Cinclocerthia* sister relationship.

The most important difference between previous studies and ours is in the phylogenetic placement of *Dumetella*. Earlier work supported a sister relationship between *Dumetella carolinensis* and *Ramphocinclus brachyurus*. Instead, we found a well-supported sister relationship between *D. carolinensis* and *Melanoptila glabrirostris* (Figs. 2–4). It should be noted that analyses of concatenated datasets in prior studies were dominated by variable sites in the mitochondrial genome, while the gene trees from single nuclear loci in those studies failed to resolve relationships among genera.

The phylogenetic placement of *Dumetella* has important implications for our understanding of the historical biogeography of this group. Prior phylogenetic hypotheses that included a sister relationship between *Dumetella* and *Ramphocinclus* require either two independent colonizations of the West Indies or a single colonization with a subsequent ‘reverse colonization’ of the mainland, at least for breeding, by an island-bound *Dumetella* ancestor (Lovette et al., 2012). Although the latter scenario has been demonstrated for birds as well as other taxa from the West Indies, and can potentially contribute to continental biodiversity (Bellemain and Ricklefs, 2008), our analyses strongly support monophyly of the resident West Indian thrashers and tremblers (i.e., excluding *Dumetella*), and thus a single colonization of the archipelago without subsequent recolonization of the mainland.

Moreover, our results provide further insight into the biogeography of temperate–tropical bird migration. Although Wallace (1874) first suggested that European birds evolved migration to escape harsh winters, most ornithologists now believe that long-distance migration evolved as tropical species expanded their breeding ranges poleward (Cox, 1968; Rappole, 1995; Somveille et al., 2018). A few recent studies have challenged that paradigm, finding considerable evidence for temperate origins of long-distance migration (Bryson et al., 2014; Winger et al., 2014; Gómez et al., 2016). Our phylogeny suggests that *Dumatella* evolved long-distance migration sometime after a split from a tropical common ancestor, providing another data point supporting the tropical origin of migration.

Species designations for resident West Indian thrashers and tremblers are validated by reciprocal monophyly and phylogenetic divergence among taxa as well as sympatry of multiple taxa on single islands (e.g., four of five species occur on Martinique and Saint Lucia). However, Lovette et al. (2012) argued that they might be oversplit at the generic level since the depth of divergence among the West Indian genera is comparable to divergence within other single genera in the family (e.g., *Mimus* or *Toxostoma*). Now, with *Dumetella* excluded from the West Indian clade, *Cinclocerthia* (Grey, 1840) would have precedence if the West Indian thrashers and tremblers were to be collapsed back into a single genus. In our view, a better alternative would be to merge *Margarops* and *Allenia* into *Cinclocerthia* but retain *Ramphocinclus*, which would capture the deepest split in the clade while collapsing the more closely related genera.

### 4.2 Species limits in Ramphocinclus *and conservation implications*

Phenotypically, the most obvious intraspecific diversity is arguably seen in *Ramphocinclus brachyurus*, which has two subspecies [*brachyurus* on Martinique and *sanctaeluciae* on Saint Lucia] that differ in plumage, ecology, and potentially calls (John, 1995; Cody, 2005). Our phylogenetic analysis shows that these subspecies are also considerably divergent genomically, with branches between the two subspecies similar in length to those that separate the other West Indian genera (Figs. 2–4). This result supports previous hypotheses that *sanctaeluciae* should be considered a separate species (Cory, 1887; Ridgway, 1907). This taxonomic change would have important implications for conservation planning: *R. brachyurus* is currently classified by the Red List as “Endangered” (IUCN, 2018). However, recognizing *R. sanctaeluciae* as specifically distinct from a Martinique endemic *R. brachyurus*, as we recommend here, would result in Red List designations of critically endangered and endangered for *brachyurus* and *sanctaeluciae*, respectively. Further characterization of genomic diversity and divergence within *R. brachyurus* based on population-level sampling is in progress (Mortensen et al., in prep).

### 4.3 Intraspecific population structure

Many West Indian bird species are distributed on multiple islands and show considerable phenotypic divergence among populations inhabiting different islands (Ricklefs and Bermingham, 2008). This pattern is also evident in the five West Indian thrashers and tremblers, each of which comprises multiple described subspecies (Cody, 2005).

The two other species for which we analyzed samples from multiple islands, *Allenia fusca* and *Cinclocerthia ruficauda*, had more shallow divergence among island populations. For *A*. *fusca*, we sampled two individuals from Dominica and one from Saint Lucia, representing two of the five described subspecies (Cody, 2005). The samples from Saint Lucia were strongly supported as monophyletic in all but the SNP-based analysis (Figs. 2–4). Divergence between populations on these two islands is also evident in mitochondrial DNA (Hunt et al., 2001), suggesting a lack of ongoing-gene flow between these populations. For *C*. *ruficauda*, we sampled populations from three islands that represent two described subspecies: *ruficauda* (Dominica) and *tremula* (Guadeloupe and Montserrat). None of our phylogenetic analyses supported reciprocal monophyly across these islands (Fig. 2–4). Nonetheless, STRUCTURE results strongly differentiated the Dominica+Guadeloupe populations from Montserrat at *K* = 2 (Fig. 5), suggesting that the former two populations have more recent shared ancestry and/or ongoing gene flow. Note that this result is not consistent with the distribution of currently recognized subspecies. Overall, our results support earlier inferences from mitochondrial sequence data that inter-island dispersal in West Indian tremblers and thrashers is effectively null in most cases, consistent with taxon cycle biogeographic models (Ricklefs and Cox, 1972; Hunt et al., 2001; Ricklefs and Bermingham, 2002; Ricklefs and Bermingham, 2008) as well as alternatives that emphasize limited dispersal in forest specialist Antillean birds (Dexter 2010).

### 4.3 Conclusions

Here we present a phylogenetic hypothesis for the West Indian thrashers and tremblers and allies that builds on previous work by using ddRAD-seq to broadly sample the nuclear genome for the first time. While most relationships were similar to those recovered in previous analyses (Hunt et al., 2001; Lovette and Rubenstein, 2007; Lovette et al., 2012), our study yields two novel and important results. First, our analyses unambiguously excluded *Dumetella* from a monophyletic group of resident West Indian species, which supports a single colonization of the archipelago without the need to posit a subsequent continental reverse colonization. Second, ours is the first phylogenetic study to sample both subspecies of the endangered *Ramphocinclus brachyurus* and assess genetic divergence between these morphologically and ecologically distinct taxa. Our results strongly support reciprocal monophyly of the two currently recognized subspecies with a level of genomic divergence typical of intergeneric comparisons in the group. This result argues for the treatment of these taxa as separate species, which would have significant conservation implications. More generally, our study enhances understanding of the evolutionary history of these birds, which represent the only avian example of adaptive radiation in the West Indies, likely owing to their early colonization of the archipelago (Ricklefs and Birmingham, 2008).

## Acknowledgments

We are grateful to the Saint Lucia Department of Forestry and Land Resources Development (Ministry of Agriculture, Natural Resources, and Co-operatives) for research permissions and logistical support and to D.C.G. Properties Limited for site access in Saint Lucia. For work in Martinique, we thank DEAL Martinique (Ministère de l’Ecologie, du Développement Durable et de l’Energie) for sponsoring permit applications; CSRPN (Conseil Scientifique Régional du Patrimoine Naturel), CRBPO (Centre de Recherches sur la Biologie des Populations d’Oiseaux), and the staff of PNRM (Parc Naturel Régional de la Martinique) for supporting our field study; and Thierry Lesales, Jean-Raphaël Gros-Désormeaux, and Georges Tayalay for logistical support and on-site assistance. Our thanks also to naturalist Stephen Lesmond and the many field technicians who helped make this work possible.

### Ethics statement

Capture and handling procedures were approved by the Tufts University Institutional Animal Care and Use Committee and follow standard best-practices as described in Fair et al. (2010), and was conducted with the requisite permits from the Governments of Saint Lucia and Martinique.

